# Indirect interactions driven by soil effects enable coexistence among competing plant species

**DOI:** 10.1101/2025.10.14.682278

**Authors:** Ezequiel Antorán, Jaime Madrigal-González, Rubén Bernardo-Madrid, Miguel Á. Fernández-Martínez, Marcelino de la Cruz, Joaquín Calatayud

## Abstract

Reciprocal effects between plants and soil have been proposed as mechanisms that promote coexistence. However, recent theoretical and empirical works have questioned their role in stabilizing coexistence within multispecies communities. In these systems, indirect interactions mediated by plant-soil feedback may play a pivotal but often overlooked role. We investigate these indirect interactions using an experimental system of two competing shrub species grown in soil conditioned by a third tree species. Tree-induced shifts in soil microbial communities, metabolites, and nutrients boost growth in the weak competitor species while reducing germination in the dominant competitor. Simulations demonstrate that these shifts in plant performance are sufficient to stabilize coexistence and to closely reproduce natural spatial distribution patterns. Our findings underscore the role of plant-soil feedback in driving indirect interactions that sustain coexistence in diverse plant communities, highlighting the importance of indirect but also positive and negative multitrophic interactions in maintaining biodiversity.

## Introduction

How can plants that use similar resources coexist? The long-standing belief that species competing for the same resources cannot stably coexist has shaped ecological theory through the lens of the competitive exclusion paradigm ^1^. Yet plants are not merely resource consumers—they actively modify soil properties and nutrient dynamics through interactions with microbial communities, mediated by processes such as nutrient uptake, litter deposition, root exudation, and allelopathy ^2–5^. These plant-driven soil modifications can, in turn, strongly influence germination, growth, and reproduction ^6–8^, shaping not only the fitness of individual plants ^9,10^ but also the performance and competitive interactions among neighbouring species ^11,12^. These reciprocal interactions—known as plant–soil feedbacks—are increasingly viewed as a potential mechanism for maintaining biodiversity by limiting conspecific dominance ^12–15^. However, despite their theoretical appeal, both empirical and theoretical evidence remain limited regarding their role in promoting species coexistence within multispecies communities ^16–18^.

Multispecies communities are characterized by pervasive indirect interactions, which occur when the dynamics between two species are influenced by the presence of a third ^19^. These interactions significantly affect plant performance ^20,21^ influencing competitive and facilitative processes, with far-reaching implications for community stability and species coexistence ^22–25^. Despite the recognized importance of indirect interactions in community dynamics, much of the current understanding of plant-soil feedback and its role in species coexistence stems from studies that isolate single-species or pairwise interactions ^8,15,26–28^. This focused approach, while informative, inherently overlooks the web of indirect effects that are intrinsic to multispecies communities, potentially contributing to the lack of consensus on the role of plant-soil feedback in promoting species coexistence in natural communities ^16,17,29^. Stabilizing indirect interactions mediated by plant-soil feedback would entail that shifts in soil biotic and abiotic characteristics caused by one species drive demographic shifts in other plant species that foster coexistence. For instance, soil modifications by a third species may reduce the fitness of dominant competitors, thereby enhancing the persistence of weaker competitors and promoting stable coexistence. Therefore, unravelling the role of plant-soil feedback in shaping multispecies coexistence requires a comprehensive investigation of its effects through the lens of indirect interactions.

To explore the role of indirect interactions mediated by plant-soil feedback in promoting multispecies coexistence, we investigated an experimental system involving three woody species. We examine how soil modifications caused by one intermediate tree species (*Quercus pyrenaica*) influence the competitive dynamics of two closely related shrub species (*Cistus ladanifer* and *Cistus laurifolius*). These shrubs are broadly distributed and dominate many shrublands across the Iberian Peninsula ^30^. Throughout much of the Mediterranean region, *C. ladanifer* often dominates, forming nearly monospecific stands. In these areas, the rare or weak competitor, *C. laurifolius* tends to occur in the presence of *Q. pyrenaica* ^31^ (Supplementary Appendix S1). *Quercus* tree species are particularly relevant in this context, as they are well known to induce strong soil modifications through root exudation and litter deposition ^32^, with extensive evidence documented for several species of the genus ^32–34^. This makes *Q. pyrenaica* an ideal focal species for studying plant–soil feedback effects. Hence, this system provides a relatively simple yet ecologically relevant context to investigate potential indirect mechanisms mediated by soil shifts in stabilizing species coexistence. Its simplicity allows us to perform controlled and detailed experiments while its broad natural distribution ensures that experimental findings can be validated with field observations. By leveraging this widely distributed but experimentally tractable system, our study aims to uncover the indirect interactions mediated by plant-soil feedback that may underpin coexistence in plant communities.

## Results and discussion

### Effects of soil conditioning by the intermediate species on germination

The majority of evidence linking species coexistence to reciprocal plant–soil interactions is based on their effects on plant growth ^13,14^. Yet, shifts in soil properties are also expected to influence critical demographic plant parameters that may facilitate the coexistence beyond growth, such as germination ^35^. Therefore, we initially investigated whether soil conditioned by the intermediate species affects the germination of the two competing species. To facilitate the link between experimental and observational evidence, we used naturally occurring soils from our study system that were undoubtedly conditioned by a single target species: *Quercus pyrenaica* (the intermediate species) and *Cistus ladanifer* (the dominant competitor). In addition, we included soils unconditioned by woody species as control soils (see Methods for details). The germination of the dominant species is significantly reduced in the influence of the intermediate species’ soil compared to dominant species’ soil (*Z* = 2.55; *P* = 0.033; Fig. 1A; Extended Data Table 1 and Supplementary Appendix S2). Comparison with the control soil reveals that the significantly reduced germination arises from a combination of increased germination in the dominant species’ soil and decreased germination in the intermediate species’ soil (Fig. 1A). In contrast, the germination of the weak competitor remains similar across all treatments (*P* > 0.05 always; Fig. 1B and Extended Data Table 1). These findings show that the intermediate species modifies the soil in ways that reduce the germination of the dominant competitor, without affecting the weak competitor. To determine whether these effects are primarily driven by soil biochemistry or living microorganisms, we compared germination responses in sterilized soils. Similar to results on live soils, germination of the dominant competitor was significantly reduced in sterilized soils previously conditioned by the intermediate species, compared to those conditioned by the dominant species (*Z =* 3.297; *P* = 0.01; Fig. 1A). This suggests that the observed effects are likely mediated by shifts in soil biochemistry, such as the accumulation of allelochemicals produced by the intermediate species ^36^. Together, these findings indicate that biochemical soil modifications induced by the intermediate species can suppress the dominant competitor, potentially promoting coexistence by reducing competitive asymmetries.

**Fig. 1.**
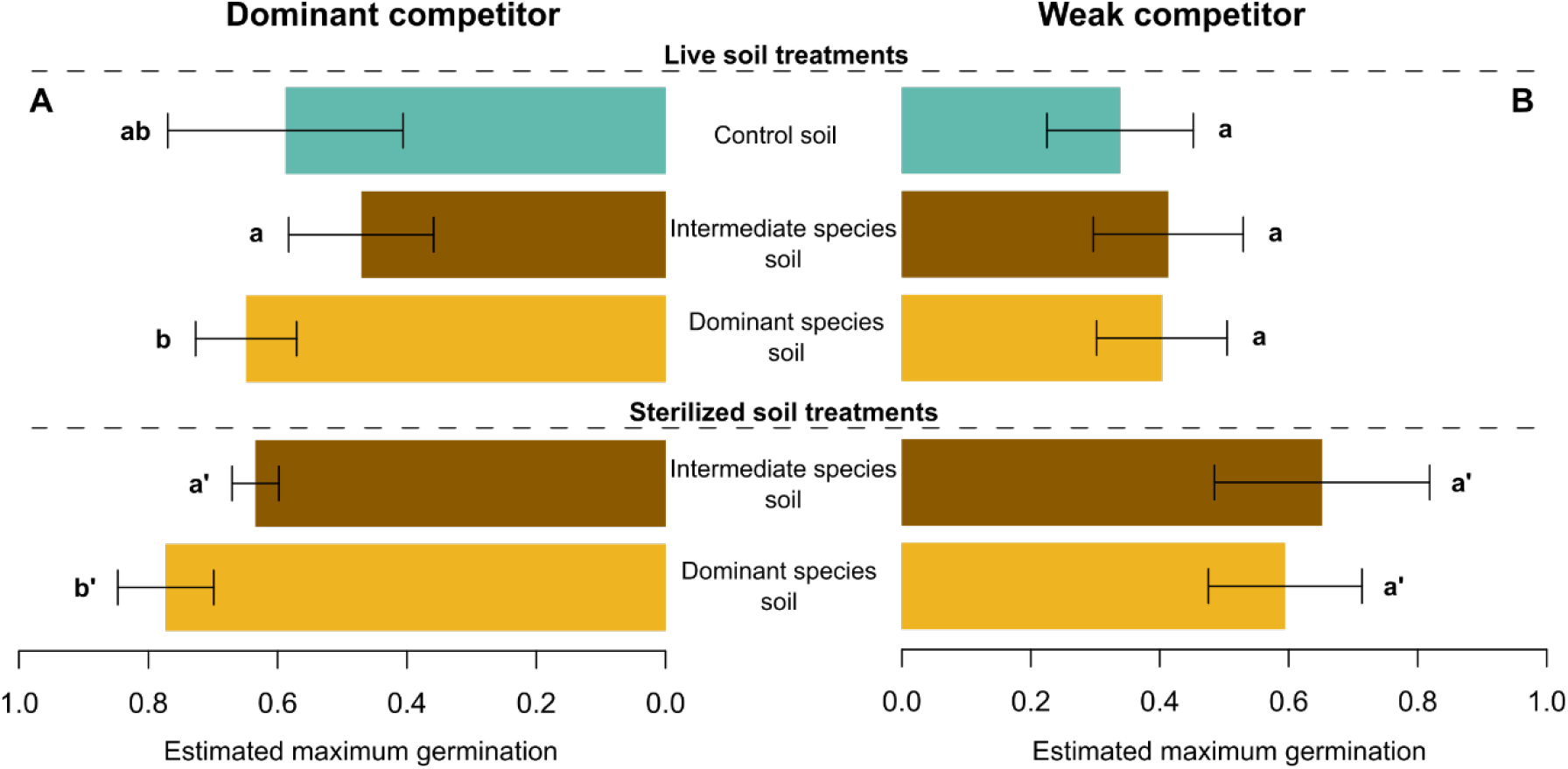
Biochemistry of soil conditioned by the intermediate species reduces the germination of the dominant competitor. The dominant competitor (*C. ladanifer*, A) exhibits significantly reduced germination in both live (top bars) and sterilized (bottom bars) soils conditioned by the intermediate species, indicating negative effects of soil biochemistry. In contrast, the weak competitor (*C. laurifolius*, B) shows no significant response to soil treatments. Bars represent the estimated maximum germination (mean and 95% confidence interval) in different soil solutions, fitted using a nonlinear logistic growth mixed model (alternative models provide similar results, Supplementary Appendix S2). Treatments were based on soils conditioned either by the intermediate species *Quercus pyrenaica* (intermediate species), the dominant species *Cistus ladanifer* (dominant species), or by none of them (control; see Methods for details). Different letters indicate statistically significant differences within sterilized and live soil treatments (P < 0.05, Extended Data Table 1).

### Effects of soil conditioning by the intermediate species on plant growth

Although the observed changes in germination may contribute to the coexistence of these competitors, the germination rates of the dominant competitor remain higher than those of the weak competitor (Fig. 1B), suggesting that additional mechanisms are involved. Plant-soil feedback is well-known to affect plant growth, which might also contribute to species performance and coexistence ^13,14^. We therefore investigated whether soil conditioned by the intermediate species affect the growth of these closely related species. In contrast to germination, no significant differences are observed in the growth of the dominant competitor across all soil treatments (*P >* 0.05 in all cases; Fig. 2A; Table S2). However, our results reveal that the growth of the weak competitor is significantly enhanced in the intermediate species’ soil compared to the control and dominant species’ soils (P < 0.01 in both cases; Fig. 2B; Extended Data Table 2). Because larger seedlings generally experience lower mortality in semiarid environments ^37^, this increase in growth likely promotes the persistence of the weak competitor under Mediterranean semiarid conditions. Taken together, these findings demonstrate that plant–soil feedback can drive demographic shifts by influencing distinct stages of plant ontogeny ^35^. More importantly, the reduced germination of the dominant species, coupled with the enhanced growth of the weak competitor in soil conditioned by the intermediate species, is expected to promote the coexistence of both competitors, underscoring how soil-mediated positive and negative interactions can support biodiversity.

**Fig. 2.**
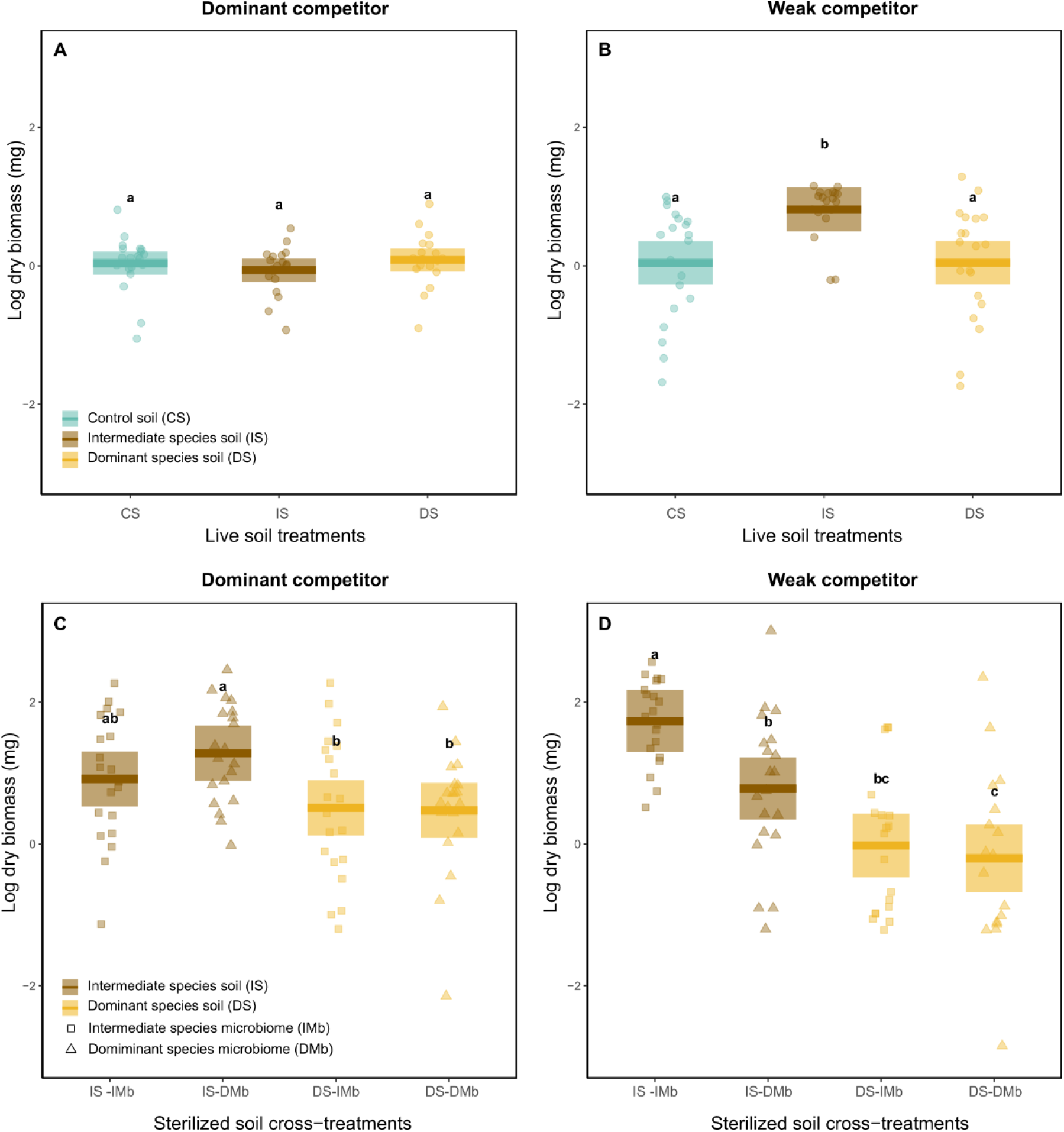
Microorganisms and biochemistry of soil conditioned by the intermediate species enhance the growth of the weak competitor. (A) Live soil treatments do not influence the growth of dominant competitor (*Cistus ladanifer*) while (B) soil conditioned by the intermediate species (*Quercus pyrenaica*) promotes the growth of the weak competitor (*C. laurifolius*). In cross-treatments using sterilized soil with microbiome inoculums, (C) The dominant competitor benefits from intermediate species’ soil only when its own soil microbiome is reintroduced. Conversely, (D) the weak competitor benefits from both soil biochemistry and the associated microbiome of the intermediate species. Box plots show log-transformed dry biomass (mg): solid bars represent means, shaded areas indicate 95% confidence intervals. Soil treatments are indicated by color, and in (C) and (D), microbiome inoculum treatments are distinguished by symbol shape. Different letters indicate statistically significant differences (*P <* 0.05, see Extended Data Table 2). CS, IS and DS refer to control, intermediate species’ and dominant species’ soils, respectively. Microbiome inoculums are abbreviated as IMb (intermediate-species) and DMb (dominant-species).

To gain deeper insights into the mechanisms behind the effects of the intermediate species on the growth of the competing shrub species, we explored whether growth changes are driven solely by soil biochemistry or also involve shifts in microbial communities. To address this, we conducted a growth experiment using sterilized soils paired with water-based microbial inoculum derived from live soils in a cross-experimental design (see Methods). In the dominant competitor, a significant enhancement in growth was observed only between plants grown in intermediate species’ soil inoculated with microorganisms of the dominant species’ soil and those grown in the dominant species’ soil, regardless of the microbial inoculum (*P <* 0.05 for both comparisons; Fig. 2C; Extended Data Table 2). Thus, the dominant competitor appears to benefit from the nutrient-rich soils conditioned by the intermediate species (see Supplementary Appendix S3) only when its native soil microorganisms are present—a scenario that does not occur under natural conditions and is consistent with the absence of effects reported above in untreated, live soils. Similarly, the weak competitor exhibits significantly enhanced growth in intermediate species’ soil when paired with microorganisms associated with the intermediate species compared with all other treatments (*P <* 0.05; Fig. 2C; Extended Data Table 2). This experimental setup, which mimics natural conditions, reveals that the enhanced growth of the weaker competitor in soils from intermediate species arises from the combined effects of shifts in soil biochemistry and microbial communities. Hence, plant growth seems to depend not only on soil chemical properties but also on specific microbial communities to which plant species are likely adapted ^38^. Importantly, our findings suggest that shifts in both soil biochemistry and microbial communities —induced by the intermediate species— are necessary to facilitate the coexistence of these closely related species.

### Experimentally based expectations align with the spatial organization of natural communities

Experimental evidence is most compelling when it directly links to patterns observed in natural communities. If the intermediate species modifies soil properties in ways that influence the germination and growth of the competing shrub species, thereby shaping their coexistence, we would expect that: (i) individuals of the intermediate species alter both biotic and abiotic soil properties at fine spatial scales, and (ii) at this scale, via soil-mediated effects, the dominant and the weak competitor species tend to segregate from and aggregate to the intermediate species, respectively. Supporting the first expectation, proximity to individuals of the intermediate species significantly shifts the composition of soil nutrients (R^2^ =7.84%; F=4.08; P < 0.01), metabolites (R^2^ =4.06%; F=2.03; P < 0.01), and bacterial (R^2^ =5.02%; F=2.54; P < 0.01) and fungal (R^2^=3.14 %; F=1.55; P < 0.01) communities. These effects are not linear but instead occur within short distances from the individuals, with compositional dissimilarities reaching their maximum at approximately 5.73, 6.28, 3.87 and 3.82 meters for nutrients, metabolites, bacterial, and fungal compositions, respectively (Fig. 3 A–D). The changes in soil composition at short distances from the tree trunk correspond to areas that remain largely covered by tree litter (Supplementary Fig. S6) throughout most of an individual’s lifespan (average age 36.6 years, Supplementary Appendix S4) and probably characterised by a denser root system. Such conditions strongly suggest that the combination of litter decomposition and root exudations are driving the pronounced modifications in soil composition observed near the tree individuals. This is consistent and aligns with extensive evidence showing that *Quercus* spp. strongly modify soil chemical, and biological properties through litter inputs and root exudations^32–34^. Our findings also support the second expectation: the abundance of the dominant competitor increases with distance from the individuals of the intermediate species (R^2^=13.6%; F=24.23; P < 0.01; Fig. 3 E), whereas the weak competitor follows the opposite trend (R^2^=3.03%; F=7.53; P < 0.01). This spatial replacement pattern is evident, with the weak competitor prevailing within approximately 3 meters from the intermediate species and the dominant competitor dominating beyond this distance (Fig. 3 E). Focusing on species cover instead of individual counts yields similar results (Supplementary Fig. S7). Although the specific soil microorganisms and chemical compounds driving these patterns remain unidentified, the fine-scale spatial structure of soil properties and species distribution closely aligns with experimental-based expectations. This supports the idea that soil modifications by intermediate species play a crucial role in promoting the coexistence of competing species.

**Fig. 3.**
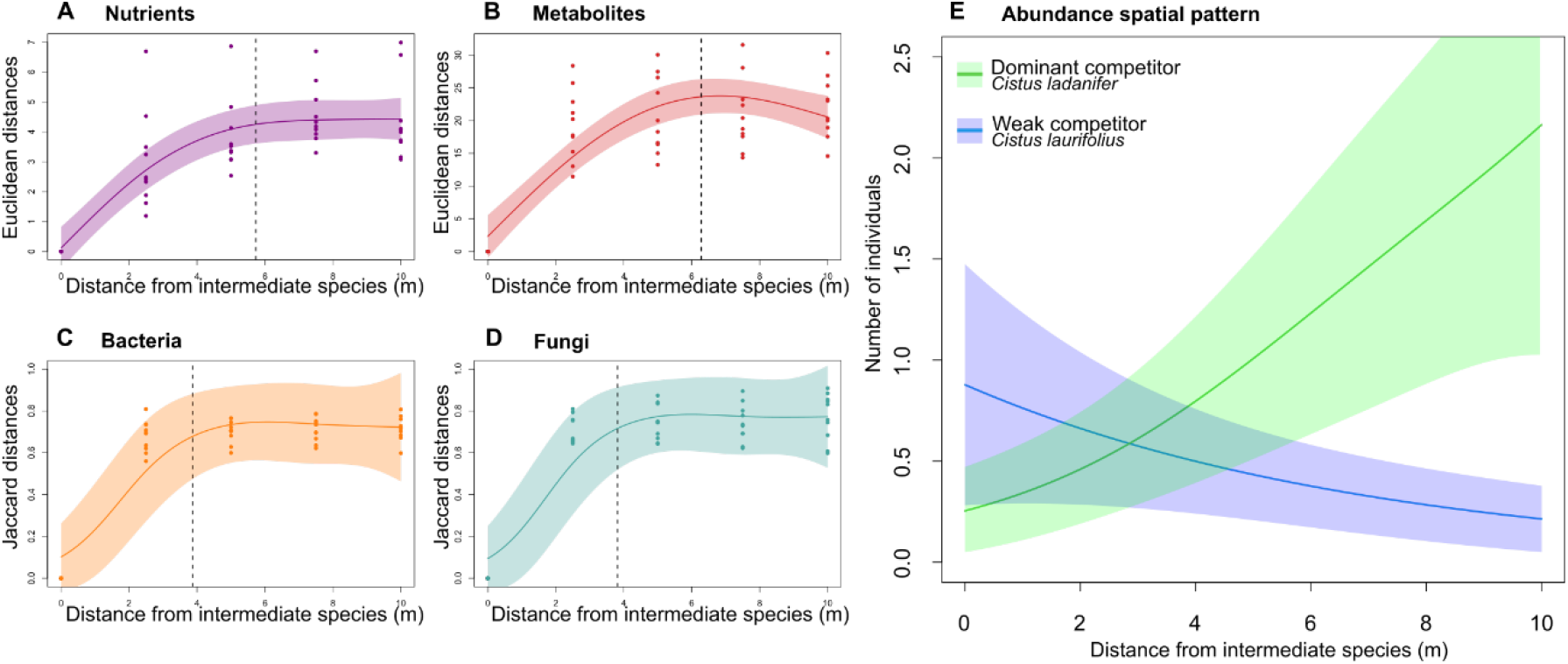
The presence of the intermediate species drastically alters the microbial communities and biochemistry of natural soils as well as the distribution patterns of the competing shrub species. The presence of the intermediate species (*Quercus pyrenaica*) significantly alters the composition of(A) nutrients (*R*^*2*^ =66.1%; *F*=85.24; *P* < 0.01), (B) metabolites (*R*^*2*^ =66.6%; *F*=69.12; *P* < 0.01), (C) bacteria (*R*^*2*^=86.3%; *F*=4.76; *P*=0.016) and (D) fungi (*R*^*2*^ =85.8%; *F*=5.17; *P=*0.012). Solid lines show predictions from a generalized additive mixed model of compositional dissimilarity between the sample collected at zero distance from individuals of the intermediate species and samples collected farther away, as a function of distance. Shaded areas represent 95% confidence intervals. The vertical dotted lines indicate the inflection point in compositional dissimilarities. (E) Predicted spatial distribution of the number of individuals in 1m^2^ plots of the dominant (*Cistus ladanifer*, green) and weak (*C. laurifolius*, blue) competitor species, relative to the distance from individuals of the intermediate species. The abundance of the weak competitor is replaced by the dominant competitor as distance from the intermediate species increases. Predictions are based on fitted generalized additive mixed models with a Poisson error distribution and a log link function.

### Simulations predict spatial patterns and stable coexistence of the competing species

Definitive evidence for the role of soil-mediated indirect interactions in stabilizing coexistence would require experimental data on all relevant demographic parameters ^35^. However, obtaining such data —e.g. seed production— in the studied woody species is largely unfeasible due to their long-life cycles and the challenges of maintaining realistic yet fully controlled experimental conditions over extended timescales. We therefore used numerical simulations parameterized with experimental results to assess whether observed changes in germination and growth are sufficient to enable the stable coexistence of both shrub species. We used experimentally derived germination data, and inferred survival probabilities from observed growth responses in conditioned soils. This inference was based on the well-supported assumption that increased growth enhances survival ^37^. Our simulations showed that the combination of measured germination probabilities and a slight increase of 0.08 in the survival probability of the weak competitor in soil conditioned by the intermediate species—compared to soil conditioned by the dominant competitor— reproduced a spatial pattern closely matching the observed distribution of both competing shrubs in natural communities (averaged *R*^*2*^*= 0*.*92*; average RMSD= 0.313; compare Fig. 4A and Fig. 3E, see also Supplementary Fig. S5 and Table S3). Under this parameterization, both species also maintained stable population sizes over time (Fig. 4B, Supplementary Table S3), establishing a link between the demographic effects of soil modifications by the intermediate species, the spatial distribution of the shrub species, and their long-term coexistence. Soil modifications by the intermediate species might also influence reproduction ^35^ or even the competitive process itself ^15^. Additionally, the intermediate species could also affect competitive dynamics through other mechanisms, such as asymmetric shading effects or differential resource use overlap with the shrub species. Nonetheless, our results suggest that only the combined effect of soil-mediated reduced germination in the dominant competitor and enhanced growth in the weaker competitor can be sufficient to enable their coexistence. We acknowledge that the dynamics of the intermediate species were not explicitly included in our framework, and therefore the long-term coexistence of the shrub species will be conditioned by the longer term demographic dynamics of the intermediate species. However, *Quercus* trees are far more long-lived than the shrubs with the lifespan of a single tree spanning several shrub generations, which likely explains the congruence between our simulations and the observed field patterns, provided that living trees are present. Overall, these results highlight the role of indirect interactions via soil effects in stabilizing species coexistence, underscoring their importance in shaping community dynamics.

**Fig. 4.**
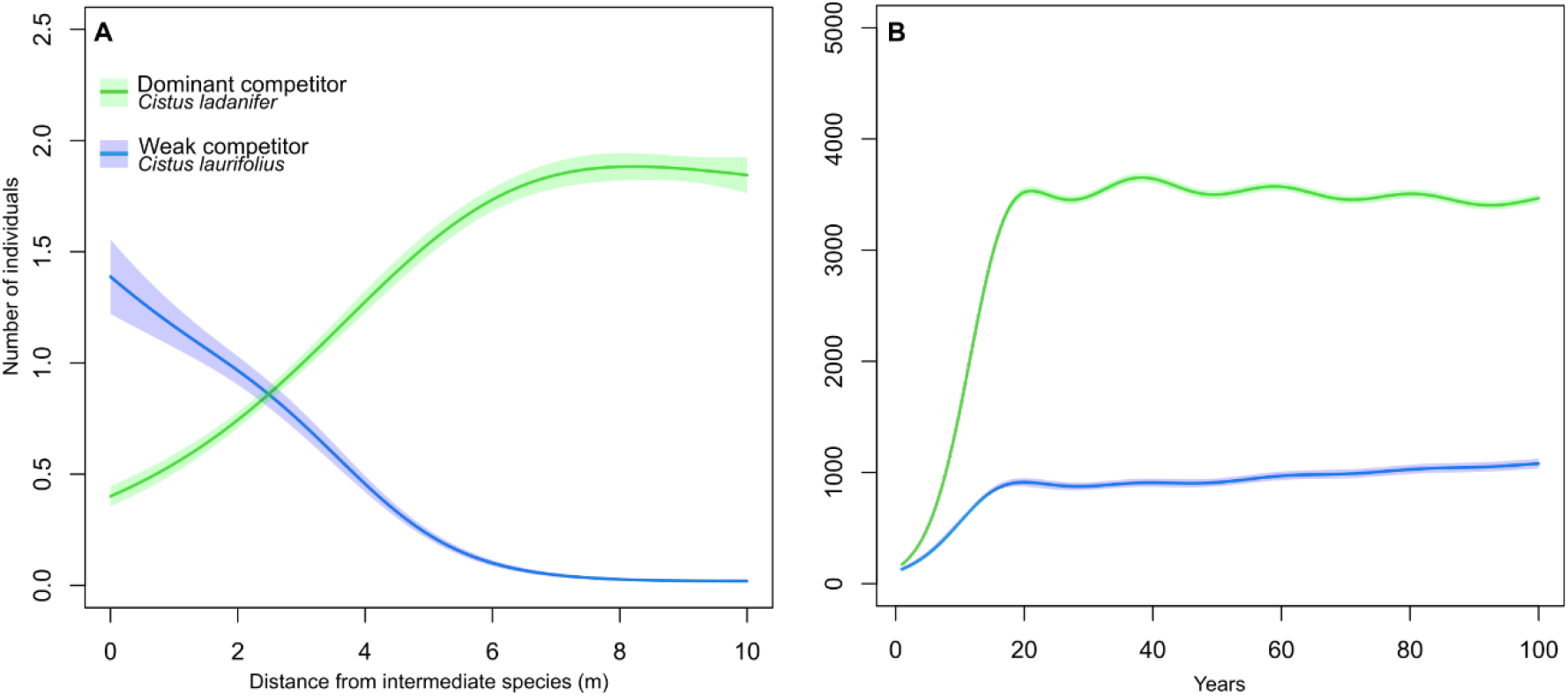
Experimentally informed simulations reproduce natural spatial patterns of competing species relative to the intermediate species, supporting also their stable coexistence. (A) Simulated spatial distribution of the number of individuals in 1m^2^ of the dominant (*Cistus ladanifer*, green) and weak (*C. laurifolius*, blue) competitor species as a function of distance from individuals of the intermediate species (*Quercus pyrenaica*). Similar to field observations (Fig. 3 E), the dominant competitor increases in abundance with distance from the intermediate species, while the weak competitor aggregates near it (*R*^*2*^=36%; *F*=619.5; *P* < 0.01 for the dominant competitor and and *R*^*2*^=22%; *F*=389.3 ; *P* < 0.01 for the weak competitor,). Model predictions (solid lines) were estimated using a generalized additive mixed model with a Poisson distribution and log link function. Shaded areas denote 95% confidence intervals. (B) Population dynamics of both competitors across the entire simulated area over 100 years of simulation. Under the observed parameterization, populations have remained stable over the last 20 years, showing no significant temporal trends for any species (*P >* 0.05). This stability supports the idea that indirect soil-mediated effects may play a key role in facilitating long-term coexistence.

## Conclusions

Our findings reveal a previously overlooked mechanism by which competing plant species can coexist in diverse ecological communities. By acting as intermediaries, certain species can modify both biotic and abiotic soil components in ways that suppress dominant competitors while facilitating weaker ones. These results highlight the need to integrate indirect and multitrophic interactions ^22,25,39^ —such as those between plants and soil microbes— into coexistence theory, moving beyond a framework centred on direct competition ^40^. Moreover, our work highlights that both positive and negative indirect interactions play a fundamental role in promoting species coexistence and sustaining biodiversity ^25,41,42^. Expanding coexistence theory to incorporate the indirect as well as positive and negative interactions will help resolve the diversity paradox, reframing species-rich communities as the expected outcome of complex ecological interactions rather than an exception ^43^. It still remains unclear which specific processes—such as root exudation, litter deposition, or the timescales over which soil modifications accumulate—are responsible for altering soil properties through the intermediate species that facilitate species coexistence. Similarly, the long-term demographic dynamics of the intermediate species and their influence on shaping the coexistence within community remain uncertain. Addressing these questions will require future long-term experimental studies spanning the full lifespan of intermediate species and ecological impacts of the species involved. Future research should also assess whether these dynamics scale to richer communities and explore how species differentially adapt to microbial communities and soil biochemistry. Addressing these questions will provide deeper insights into the eco-evolutionary pathways through which indirect plant-soil feedback mediate coexistence. More broadly, our findings reinforce the view that the maintenance of biodiversity is shaped not only from competition but from a web of interacting forces, where soil-mediated positive and negative indirect effects can play a central role in stabilizing species-rich communities.

## Methods

### Study system

We used two closely related mediterranean shrub species (*Cistus ladanifer* and *Cistus laurifolius*) and a *Quercus* tree (*Quercus pyrenaica*) as our experimental system. *C. ladanifer*, is widely distributed across the southern and western regions of the Iberian Peninsula ^44^, whereas *C. laurifolius* predominantly occupies the northeastern half ^45^. A broad contact zone exists between these species, with the “Sistema Central” mountain range representing the primary area of overlap. Within this contact zone, both two species are generally segregated by altitude: *C. ladanifer* dominates the lowlands, while *C. laurifolius* is more prevalent at higher elevations ^46^. However, *C. laurifolius* can occasionally be found in lowland areas where *C. ladanifer* is dominant, but normally in the presence of *Q. pyrenaica* (^31^; Supplementary Appendix S1). We focused on these lowland areas, and based on species’ abundances, we define *C. ladanifer* as the dominant competitor, given its high abundance, and *C. laurifolius* as the weak competitor, as it is much less frequent in these areas. *Quercus pyrenaica* is considered the intermediate species, as its presence is associated with the occurrence of *C. laurifolius* in lowlands otherwise dominated by *C. ladanifer* (Supplementary Appendix S1). We specifically examine how the two competing shrub species respond to soil modifications associated with the presence of the intermediate species. Given their close phylogenetic relationship and high resource overlap, *C. ladanifer* and *C. laurifolius* are expected to engage in strong interspecific competition ^47^.

### Study area, soil and seed collection and treatments

The study area is located at the south of the Sistema Central mountain range in central Spain, at an elevation of 940-1050 m a.s.l. (40°26′ 46′′N, 3°42′ 36′′W). It features a continental Mediterranean climate characterized by cold winters and mild, dry summers. The landscape is dominated by extensive shrublands of *C. ladanifer*, interspersed with patches of *Q. pyrenaica* and other deciduous species.

Fruits (capsules) of wild specimens of both *Cistus* species were collected between September and October 2022 from 10 individuals per species, randomly selected across the study area. After collection, seeds were separated from the fruits, air-dried under ambient conditions, and pooled by species, with no distinction made between individual plants. Immediately before experiments, seeds were exposed to a heating treatment, 100ºC during 5 minutes, to promote germination rates ^48^.

Soil samples were collected from the study area, with sampling locations carefully selected based on vegetation type and the presumed soil conditioning effects of target species. Sampling encompassed three soils categories: (i) Dominant species’ soil was obtained from ten randomly selected places where only *C. ladanifer* was present, ensuring non-external influence from other woody species by maintaining a minimum distance of 50 m from the nearest *Q. pyrenaica* and of 20 m to other woody species. (ii) Intermediate species’ soil was collected within 50 cm of the base of ten randomly selected *Q. pyrenaica* individuals, with no other woody species within at least 20 m; and (iii) Control soil, sampled from an open area with no woody vegetation present for over 50 years, as verified through historical orthophotos and field inspection. Since the weak competitor, *C. laurifolius*, is always associated with *Q. pyrenaica* in the study area, it was not possible to obtain soils exclusively conditioned by *C. laurifolius* with full confidence. All soils were collected using ethanol-sterilized shovels to avoid contamination. Soil was extracted from an area of approximately 0.5 m^2^, at a depth of 3–20 cm, while discarding the topmost surface soil layer. Soil samples were pooled together by soil type and immediately sieved using a sterilized 60 mm mesh to remove rocks and coarse roots, ensuring a thorough mixing and standardization. Fresh live soils were collected in April 2023 and stored at 4ºC until experiments were carried out.

To distinguish the effects of the chemical and biotic components of soil, we sterilized 15 L of each soil type with gamma irradiation (25 kGy) at Aragogamm S.A. (Barcelona, Spain). For each soil type (dominant and intermediate species soils) and treatment (live soil and sterilized), we prepared an aqueous inoculum. Live soil inoculum was used in germination and in the cross-treatments of the growth experiments, while sterilized soil inoculum was only used in germination experiments. Soil aqueous inoculum were achieved by mixing 700 g of sieved soil with 1.5 L of demineralized water ^49^. The mixture was manually homogenized in a sterile container, passed through a 200 μm sterilized mesh sieve, and collected in a sterilized glass jar. Inoculum were prepared independently in sterile recipients on the same day as inoculation to prevent contamination.

### Germination experimental design and analysis

To investigate soil effects on the germination of both *Cistus* species we conducted a controlled germination experiment using five soil treatments. Each treatment involved a soil aqueous inoculum derived from different soil types: (i) soil conditioned by the intermediate species (*Quercus pyrenaica*), (ii) soil conditioned by the dominant species (*Cistus ladanifer*), (iii) unconditioned control soil, (iv) sterilized soil conditioned by the intermediate species, and (v) sterilized soil conditioned by the dominant species. To germinate the seeds, Petri dishes were used ensuring stable humidity and preventing airborne contamination, also a sterile filter paper disc was placed inside each Petri dish to ensure a balance between hydration and aeration for the seeds ^48^. For each treatment, 10 mL of the corresponding inoculum were pipetted onto the Petri dishes before seed placement. After inoculation, 20 seeds of a single *Cistus* species were evenly distributed on the moistened filter paper and spaced apart to facilitate individual observation. We established five replicate dishes per treatment. A total of 1,000 seeds were used in the experiment (20 seeds × 5 Petri dishes × 5 treatments × 2 species). Petri dishes were placed in a phytotron under controlled conditions, maintaining a cycle of 20°C of temperature during 14 hours of light and 16°C during 10 hours of darkness. The experiment lasted 30 days between April to May of 2023, during which Petri dishes were examined every three days to record the number of germinated seeds, i.e., which the emerging radicle visible to the naked eye. To prevent evaporation and maintain sufficient humidity for germination, distilled water was replenished as needed in each dish. Additionally, Petri dishes were fully randomized within the phytotron at each observation interval to eliminate possibles positional effects.

We fitted a three-parameter logistic growth model to describe the cumulative germination proportion over time for each species and soil treatment:

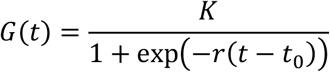

where *G*(t) is the proportion germinated at time t, *K* is the maximum germination proportion, *r* is the growth rate, and t_0_ is the time at which 50% of *K* is reached (inflection point). To account for non-independence of repeated observations within Petri dishes, we fitted a nonlinear mixed-effects model, including Petri dish identity as a random term. These models capture well the sigmoidal pattern typical of seed germination (median pseudo- *R*^*2*^*=*83%; bootstrapped 95% confidence interval: 72.10% - 88.35%, Supplementary Table S2, Fig. S2) and leverage the full temporal resolution of our data, providing a robust estimate of germination dynamics ^50^. Nevertheless, results were consistent when using an alternative model (Supplementary Appendix S2; Table S1 and S2). We focused on the *K* parameter —representing the maximum germination— and performed pairwise statistical comparisons between treatments using fitted *K* values and their standard errors to conduct Z-tests ^51^. To account for multiple comparisons, we applied a false discovery rate correction to the p-values ^52^. It should be noted that direct comparisons between sterilized and live soil treatments were not performed, as gamma sterilization altered soil nutrients ^53^. Models were fitted using the nlme package ^54^ in R ^55^.

### Growth experimental design and analysis

We conducted common garden experiments to explore the effects of soil on plant growth. The experiment consisted of two steps. First, we tested the influence of three provenances of live soils on growth: i) intermediate species’, ii) dominant species’, and iii) control soils. Second, we designed a cross-treatment experiment to disentangle the individual and combined effects of soil biochemistry and living microorganisms. The cross-treatment experiment consisted in a total of four treatments, in which sterilized soils of the intermediate and dominant species were supplemented with aqueous live-soil inoculums of the intermediate and dominant species to restore microbiological activity: i) sterilized soil of the intermediate species + aqueous inoculum of the intermediate species; ii) sterilized soil of the intermediate species + aqueous inoculum of the dominant species; iii) sterilized soil of the dominant species + aqueous inoculum of the intermediate species; iv) sterilized soil of the dominant species + aqueous inoculum of the dominant species.

For the common garden experiments, pre-heated seeds were sown into 3.5 cm × 3.5 cm (58 cc) pots using sterile forceps, which were sterilized between treatments. Each pot contained five seeds of the same species to ensure germination success. After germination, one randomly selected seedling per pot was retained for further growth. In the cross-treatment, microbiome inoculums were introduced immediately before sowing by adding 58 mL of the corresponding soil aqueous inoculum to each sterilized soil pot. We replicated each treatment 20 times for each species. A total of 280 individuals (20 individuals × 7 treatments × 2 species) were grown under controlled greenhouse conditions, with an average temperature of 22°C and 46 g/m^3^ humidity. Plants were irrigated three times daily for two minutes using a nebulizer, and supplemental LED lighting (230l μmol m^−2^ s^−1^) was provided for a 14-hour light and 10-hour dark cycle. Pots were periodically randomized to eliminate positional effects. The experiment was conducted from beginning of April to end of May, with a growth period lasting 60 days. At the end of the experiment, the aboveground part of the plants was harvested by carefully removing from the pots. Fresh biomass was then dried at 60°C for 48 hours ^56^ and weighed using a precision balance (accuracy, 1 μg; Mettler Toledo) to obtain aboveground biomass measurements. To estimate the average growth per treatment, we fitted linear regression models to the log-transformed dry biomass (mg) as a function of soil treatment, and compared marginal means applying a Tukey correction for multiple comparisons as implemented in the R package emmeans ^57^.

### Field observations

We further assessed whether the experimental results align with the spatial distribution patterns of the competing shrub species relative to the intermediate species observed in natural communities. For this, we performed vegetation inventories around *Q. pyrenaica* individuals within a ca. 2-hectare area of the study area. Ten adult trees (average age 36.6 years, average DBH 45.9 cm, Supplementary Appendix S4) were randomly selected, ensuring a minimum distance of 20 meters from other *Q. pyrenaica* individuals to prevent overlapping zones of influence. From the base of each selected tree, we established a 10 × 1 m transect in a randomly assigned direction. Each transect was subdivided into ten 1 × 1 m sections. Within each section, we recorded the number of individuals and cover of each *Cistus* species. To assess how both *Cistus* species abundances varied with distance from *Q. pyrenaica*, we modelled the recorded number of individuals and cover of each species along the transects sections. We used a generalized additive mixed model (GAMM), assuming a Poisson error distribution and a log link function and using tree identity as a random term. GAMMs were fitted using the mgcv package ^58^ in the R software.

Moreover, to explore the influence of the intermediate species on soil nutrients, metabolites composition, and microbial communities, we collected soil samples along the same transects. Sampling was conducted at five points along each transect, beginning at the base of the trunk (0 meters) and extending up to 10 meters, with a separation of 2.5 meters. At each sampling point, using a 5 cm diameter × 20 cm steel cores, three soil cores were extracted, contiguous but spaced 20 cm apart to ensure a representative sample of the soil. Cores were taken from a depth of 3–20 cm discarding the topmost surface organic layer. Soil samples from each sampling point were pooled into a single sterile, airtight bag, immediately refrigerated upon collection, sieved, and subsequently stored at −20°C until further analysis. To prevent cross-contamination, all sampling equipment, including steal corers and the mallet, was sterilized between samples using ethanol cleansing followed by controlled fire exposure. Additionally, we wore sterile gloves and masks throughout the process to minimize external and cross contamination. The collected soil samples were subsequently analysed to determine their chemical properties, metabolomic composition, and microbial (bacteria and fungi) community composition.

### Soil nutrient analyses

Soil samples were analysed at NutriLab (University of Rey Juan Carlos, Spain) to assess nutrient content and soil properties for carbon, total nitrogen, total phosphorus, pH, conductivity, nitrates, phosphates, and ammonium. Samples were dried at 65 °C, ground, and sieved (2 mm mesh). Carbon content was determined by oxidation with potassium dichromate, followed by spectrophotometric measurement at 600 nm. Total nitrogen was quantified using Kjeldahl digestion with sulfuric acid and a catalyst composed of 1.5% CuSO_4_·5H_2_O and 2% Se, with absorbance measured at 630 nm. Total phosphorus was determined after UV digestion, forming a molybdate complex detected at 880 nm. Nitrates were reduced using a Cd-Cu column and analysed via the Griess reaction at 540 nm, while phosphates were quantified by reaction with ammonium heptamolybdate and antimony potassium tartrate at 880 nm. Ammonium was determined using the Berthelot reaction, producing a green complex measured at 630 nm. Soil pH was measured potentiometrically in a 1:2 soil-water suspension, and conductivity was recorded using a conductivity meter.

### Microbiome analysis

DNA was extracted from soil samples using the HigherPurity™ Soil DNA Isolation Kit (Canvax Reagents SL, Valladolid, Spain), following the manufacturer’s instructions, including a negative control (DNA extraction blank). The extracted DNA was resuspended in 100 μL of elution buffer and its concentration was quantified using the Qubit High Sensitivity dsDNA Assay (Thermo Fisher Scientific). DNA metabarcoding library preparation and sequencing of bacterial 16S rRNA and fungal ITS2 gene regions were conducted by AllGenetics & Biology SL, employing Bakt_341F and Bakt_805R primers ^59^ and gITS7 ^60^ and ITS4 primers ^61^, respectively. Equimolar pooling of libraries was performed before sequencing on an Illumina NovaSeq PE250 platform. Raw sequencing reads were processed with the DADA2 pipeline in R software ^62^ to generate amplicon sequence variants (ASVs), following a custom modification to better suit reads characteristics.

### Metabolomic analysis

Soil samples were lyophilized before metabolite extraction. For each sample, 100 mg of dry soil were extracted with 1 mL of 80% (v/v) methanol (–20 °C), incubated at 4 °C for 10 min (2,000 rpm), sonicated for 15 min in ice-cold water, and centrifuged at 14,800 rpm for 15 min at 4 °C. The supernatant (550 μL) was dried in a vacuum concentrator (SPD1030, Thermo Fisher) and stored at –80 °C. Prior to analysis, samples were resuspended in 55 μL of acetonitrile:water (80:20, v/v) containing internal standards. Samples were analysed on a Vanquish Horizon UHPLC coupled to an Orbitrap Exploris 120 mass spectrometer (Thermo Fisher), using a BEH Amide column. Chromatographic separation lasted 17 min using a gradient with ammonium formate and formic acid modifiers. An untargeted metabolomics approach was applied to detect a broad spectrum of small polar and semi-polar metabolites present in the soil. Data were acquired in positive mode with data-dependent MS/MS (Top4). Raw files were processed with MS-DIAL v4.9.22 using *in-house* RT–m/z libraries, LipidBlast, and NIST MS/MS databases. Metabolite annotation followed the confidence criteria proposed by the Metabolomics Standards Initiative ^63^. Duplicated metabolites, adducts and isotopes were removed. Data were normalized using the Systematic Error Removal using Random Forest (SERRF) method, with pooled mixtures used as reference samples ^64^. All analyses were conducted at the Laboratory of Foodomics, Institute of Food Science Research (CIAL, CSIC), Madrid, Spain.

### Statistical Analysis

To assess differences in microbial communities, nutrients and metabolites in relation to tree proximity, we first calculated compositional distances across all soil samples. For microbial communities, we computed Jaccard distances separately for bacterial and fungal ASVs, choosing this presence/absence-based dissimilarity index to mitigate issues associated with compositional data of ASV counts ^65^. To reduce noise from rare occurrences, we excluded taxa present in fewer than five samples, though including these taxa yielded similar results (Supplementary Appendix S5). For nutrients and metabolite compositional differences, we used Euclidean distances. We conducted permutational multivariate analysis of variance on compositional distances as a function of spatial distance, as implemented in the vegan R package ^66^, incorporating tree identity as a stratification factor. Additionally, we examined where the strongest compositional differences occurred in relation to tree proximity. Specifically, we compared the compositional distances between soil sampled directly beneath tree individuals (0 m) and all the other samples. In this case, we fitted GAMMs of compositional distances as a function of spatial distances, including tree identity as a random factor. We identified turning points as locations where the first derivative of the fitted curve crossed zero.

### Simulations

We further investigate whether the experimentally observed changes in species performance are sufficient to enable the stable coexistence of both competing shrub species. To address this, we developed an individual-based simulation model in which individuals of both competing shrub species are ecologically identical, except for the demographic parameters derived from our experiments. The model consists of a lattice of 50 × 50 grid cells, each representing one square meter. We randomly placed 20 individuals of *Q. pyrenaica* across the lattice and assumed that their soil-conditioning effects extended homogeneously in a circular area with a four-meter radius around each individual. This threshold was based on observed average changes in soil microbial communities, nutrient levels, and metabolite composition (Fig. 3 A-D). At the start of the simulation, we also placed randomly 100 individuals of each competing species. In grid cells conditioned by *Q. pyrenaica*, the two competitor species performance (i.e., germination and survival probabilities, see below) was set according to estimates from experimental results on *Q. pyrenaica*-conditioned soils. In all other cells, performance was set according to probability estimates from experimental results on *C. ladanifer-*conditioned soil.

During simulations, germination probabilities were drawn from uniform distributions defined by the 95% confidence intervals estimated in the experiments for each species and soil type (Supplementary Table S2). To further include the effects of soil conditions on plant growth, we assumed that plant size influence survival probabilities (Ps) according to Andivi*a* et al., (*2*021) ^37^, following the logistic equation:

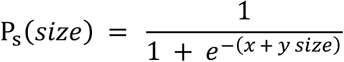

where x represents the log-odds of basal survival probability, set at 1.386 (equivalent to a survival probability of 0.8) to approximate the demographic dynamics of these *Cistus* species (Supplementary Appendix S6). The parameter *y* modulates the effect of plant size on survival probability: *y=*0 results in a constant survival probability of 0.8 regardless of size, while higher *y* values increase the effect of plant size on survival. As the actual relationship between plant size and survival remains unknown, we conducted simulations across 21 values of *y* from 0 to 2, with 0.1 increments. For each competing species and soil type, survival probabilities were drawn from a uniform distribution bounded by the probabilities estimated from the above logistic equation and plant sizes within the 95% mean confidence intervals obtained experimentally (Supplementary Table S3). Remaining model specifications were identical for individuals of both species and are detailed in Supplementary Appendix S6. For each y value we run 25 simulations, each spanning 100 iterations (representing years).

At the end of each simulation, we assessed (i) whether both competitor species persisted and maintained stable population sizes (Supplementary Appendix S6), and (ii) the degree of correspondence between simulated and field-observed abundance distributions. Population stability was evaluated by first fitting GAMMs of the total number of individuals in the lattice as a function of time for each species, using a Poisson error distribution with a log link function and including simulation replicate as a random term. We then tested for significant temporal trends over the last 20 years by applying a linear regression to the predicted abundances from the previous GAMM model as a function of time. A lack of significant abundance-time relationships (*P >* 0.05) for both competing species was interpreted as evidence of population stability (Supplementary Table S3). To assess the correspondence between observed and simulated abundance distributions, we sampled simulated abundance similarly to the field-based observations described above. To mimic the observational data-gathering protocol, we recorded the number of individuals per competing shrub species along transects of 10 grid cells extending from *Q. pyrenaica* individuals. To select a transect for each *Q. pyrenaica* individual, we established radial transects at 20° intervals and selected those that did not overlap with the soil influence of other *Q. pyrenaica* individuals. For each y-value (i.e., the parameter modulating the effect of plant size on survival probability), simulated data were analysed using GAMMs to predict species density as a function of distance from *Q. pyrenaica*, with tree individual identity nested within simulation replicate included as a random term. A Poisson error distribution with a log link function was assumed. We compared simulated and observed abundance predictions using the mean square root deviation, averaged across both competing species. We selected the *y*-value that provided the most accurate prediction of the spatial pattern of both species.

## Acknowledgements

This research was supported by the projects PID2020-114851GA-I00 (UNIPER) (Spanish Ministry of Science and Innovation) and RARABUN 2022/00156/001 (Comunidad de Madrid Government), granted to JC. EA was supported by a predoctoral fellowship (grant PRE2021-097730). JC was also supported by the Ramón y Cajal programme (RYC2021-034013-I), funded by the Spanish Ministry of Science and Innovation.

## Author contributions

EA and JC conceived the ideas and designed the experiments, with inputs from JMG. EA and JC conducted the fieldwork and experiments. EA carried out the analyses with assistance from JC and input from RBM, MAFM, and MC. EA led the writing of the manuscript with support from JC. All authors read and commented on the manuscript.

## Competing interest

The authors declare no competing interests.

## Data and code availability

All data and code used to generate the results presented in this study will be made publicly available upon acceptance of the manuscript.

---

**Extended Data Table 1.** Pairwise comparisons of the fitted K parameter for germination probability using Z-test with FDR correction. Comparisons were conducted for *Cistus ladanifer* and *Cistus laurifolius* germination under sterilized and live soil treatments: (CS) control soil, (DS) dominant-species (*C. ladanifer*) soil, (IS) intermediate-species (*Quercus pyrenaica*) soil, (SDS) sterilized dominant-species soil and (SIS) sterilized intermediate-species soil. The Z-stat refers to the Z-statistic from the pairwise comparisons of model parameters. The *P* (FDR) shows the p-values adjusted using the FDR correction to control for multiple testing.

| Species | Treatment | vs | Treatment | Z-stat | *P*(FDR) |
| --- | --- | --- | --- | --- | --- |
| *C. ladanifer* | DS | - | CS | 0.603 | 0.547 |
|  | DS | - | IS | 2.555 | 0.033 |
|  | CS | - | IS | 1.075 | 0.423 |
|  | SDS | - | SIS | 3.297 | 0.001 |
| *C. laurifolius* | DS | - | CS | 0.838 | 0.603 |
|  | DS | - | IS | -0.125 | 0.900 |
|  | CS | - | IS | -0.903 | 0.603 |
|  | SDS | - | SIS | -0.549 | 0.583 |

**Extended Data Table 2.** Post hoc Tukey comparisons from linear models assessing soil treatment effects on aboveground dry biomass of *C*.*ladanifer* and *C. laurifolius*. To denominate the treatments we use: (CS) Control soil; (IS) Intermediate-species (*Quercus pyrenaica*) soil and (DS) Dominant-species (*Cistus ladanifer*) soil, (IMb) Intermediate-species microbiome and (DMb) for Dominant-species microbiome.

|  |  | Treatm<br>ent | v<br>s | Treatment | Estimate | SE | T-ratio | p-value |
| --- | --- | --- | --- | --- | --- | --- | --- | --- |
| *C. ladanifer* | Live soil treatments | IS | - | CS | -0.100 | 0.118 | -0.851 | 0.673 |
|  |  | IS | - | DS | -0.147 | 0.118 | -1.251 | 0.429 |
|  |  | CS | - | DS | -0.047 | 0.118 | -0.400 | 0.916 |
|  | Sterilized soil<br>cross-treatments | IS-IMb | - | IS-DMb | -0.364 | 0.276 | -1.322 | 0.552 |
|  |  | IS-IMb | - | DS-IMb | 0.406 | 0.276 | 1.471 | 0.460 |
|  |  | IS-IMb | - | DS-DMb | 0.442 | 0.276 | 1.604 | 0.383 |
|  |  | IS-DMb | - | DS-IMb | 0.770 | 0.276 | 2.793 | 0.033 |
|  |  | IS-DMb | - | DS-DMb | 0.807 | 0.276 | 2.926 | 0.023 |
|  |  | DS-IMb | - | DS-DMb | 0.037 | 0.276 | 0.133 | 0.999 |
| *C. laurifolius* | Live soil treatments | IS | - | CS | 0.773 | 0.223 | 3.466 | <0.01 |
|  |  | IS | - | DS | 0.771 | 0.223 | 3.456 | <0.01 |
|  |  | CS | - | DS | -0.002 | 0.223 | -0.01 | 1.000 |
|  | Sterilized soil<br>cross-treatments | IS-IMb | - | IS-DMb | 0.951 | 0.311 | 3.059 | 0.016 |
|  |  | IS-IMb | - | DS-IMb | 1.755 | 0.315 | 5.571 | <0.01 |
|  |  | IS-IMb | - | DS-DMb | 1.933 | 0.324 | 5.961 | <0.01 |
|  |  | IS-DMb | - | DS-IMb | 0.804 | 0.315 | 2.551 | 0.06 |
|  |  | IS-DMb | - | DS-DMb | 0.982 | 0.324 | 3.029 | 0.018 |
|  |  | DS-IMb | - | DS-DMb | 0.179 | 0.328 | 0.545 | 0.948 |

## Notes

**Statement of authorship:** EA and JC conceived the ideas and designed the experiments, with inputs from JMG. EA and JC conducted fieldwork and experiments. EA carried out the analyses with assistance from JC and input from RBM, MAFM, and MC. EA led the writing of the manuscript with support from JC. All authors read and commented on the manuscript.

### Competing Interest Statement

The authors have declared no competing interest.

