## Supplementary information for "Indirect interactions driven by soil effects enable coexistence among competing plant species"

### Appendix S1. Landscape vegetation inventory

The study area is characterized by extensive shrublands interspersed with patches of deciduous forest. To accurately characterize the vegetation composition across the landscape, we randomly selected 15 sampling points within each of the two dominant vegetation types—shrubland and deciduous forest patches. At each point, we established a 5 m diameter plot and recorded the cover of all woody species present. Shrubland communities (Fig. S1 A) are overwhelmingly dominated by *C. ladanifer*, which accounts for over 85% of the total vegetation cover among the 10 species recorded. In contrast, (Fig. S1 B) forest patches are primarily dominated by *C. laurifolius* and *Q. pyrenaica* (48.7% 15.4% of cover respectively), with a total of 15 species identified in these areas. This contrast highlights the near-monospecific nature of shrubland vegetation, predominantly dominated by *C. ladanifer*, whereas forest patches support more woody species composition, in where *C. laurifolius* commonly associates with *Q. pyrenaica*.

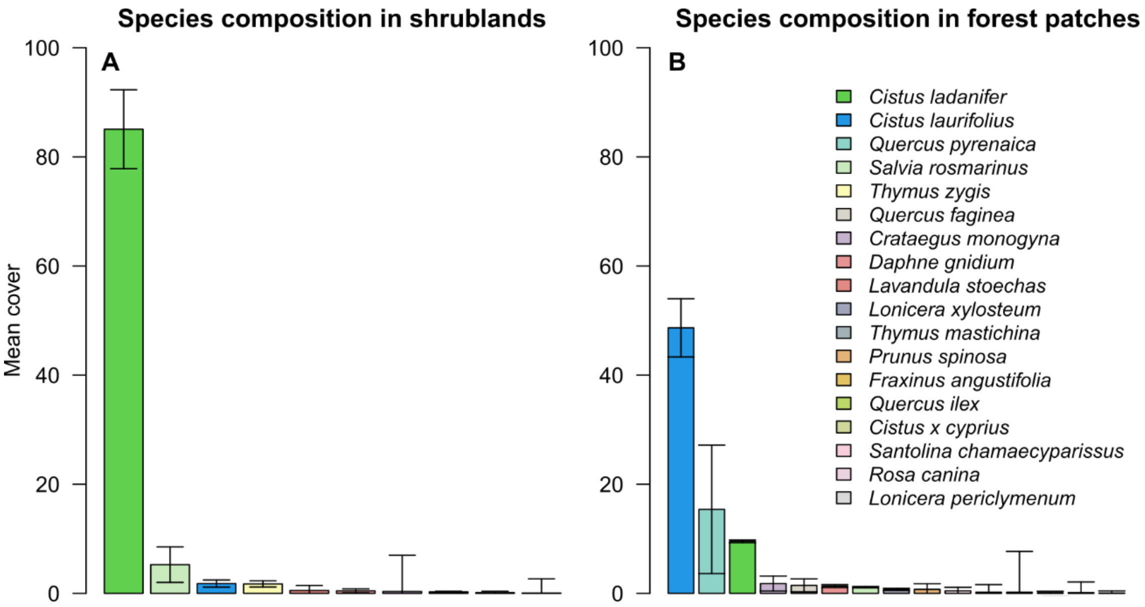

**Fig. S1. Vegetation composition in shrublands and deciduous forest patches.** *C. ladanifer* dominates shrublands, while *C. laurifolius* is commonly associated with *Q. pyrenaica* in deciduous forest patches. Bars represent the mean cover of woody species and the 95% confidence interval. (A) Shrublands are nearly monospecific, predominantly dominated by *C. ladanifer* (green). In contrast, (B) deciduous forest patches exhibit greater species diversity.

**Appendix S2. Sensitivity analysis germination models**

To assess differences in germination across soil treatments, we initially employed a logistic growth model given that seed germination typically follows a sigmoidal pattern. However, to evaluate the robustness of our findings, we also tested the Gompertz model (Gompertz, 1997) as an alternative nonlinear regression model. These model, like the logistic growth model, describe sigmoidal growth but with varying flexibility in their shape parameters. Our results remained qualitatively and quantitatively similar (Table S1 and S2) indicating that the model choice does not alter the interpretation of germination results.

**Table S1**. Pairwise comparisons of the fitted K parameter for germination probability using Z-test with FDR correction for the Logistic growth and Gompertz models. Comparisons were conducted for *Cistus ladanifer* and *Cistus laurifolius* germination under live soil and sterilized soil treatments: (CS) control soil, (DS) dominant specie' (*C. ladanifer*) soil, (IS) intermediate species' (*Quercus pyrenaica*) soil, (SDS) sterilized dominant species' soil and (SIS) sterilized intermediate species' soil. The Z-stat refers to the Z-statistic from the pairwise comparisons of model parameters. The *P* (FDR) shows the p-values adjusted using the FDR correction to control for multiple testing.

| | | | | Logistic growth model | | Gompertz model | |
|---|---|---|---|---|---|---|---|
| Species | Treatment | vs | Treatment | Z-stat | *P*(FDR) | Z-stat | *P*(FDR) |
| *C. ladanifer* | DS | - | CS | 0.603 | 0.547 | 0.656 | 0.512 |
| | DS | - | IS | 2.555 | 0.033 | 2.612 | 0.027 |
| | CS | - | IS | 1.075 | 0.423 | 1.070 | 0.428 |
| | SDS | - | SIS | 3.297 | 0.001 | 3.340 | 0.001 |
| *C. laurifolius* | DS | - | CS | 0.838 | 0.603 | 0.815 | 0.639 |
| | DS | - | IS | -0.125 | 0.900 | -0.019 | 0.985 |
| | CS | - | IS | -0.903 | 0.603 | -0.796 | 0.639 |
| | SDS | - | SIS | -0.549 | 0.583 | -0.535 | 0.593 |

**Table S2**. Results of nonlinear regression models (logistic growth and Gompertz model) for germination of *C ladanifer* and *C lauriflolius*, with all treatments: (CS) control soil, (DS) dominant-species (*C. ladanifer*) soil, (IS) intermediate-species (*Quercus pyrenaica*) soil, (SDS) sterilized dominant-species soil and (SIS)sterilized intermediate-species soil. The table reports the coefficient of determination ($R^2$), the estimated maximum germination proportion (K), the standard error (SE) of K, and the 95% confidence intervals (95% CI). The 95% CIs estimated from the logistic model were used to define the range of germination probabilities in the simulations.

| | | Logistic growth model | | | | Gompertz model | | | |
|---|---|---|---|---|---|---|---|---|---|
| Species | Treatment | $R^2$ | K | SE | 95% CI | $R^2$ | K | SE | 95% CI |
| C. ladanifer | CS | 0.536 | 0.588 | 0.182 | 0.406 - 0.770 | 0.543 | 0.594 | 0.184 | 0.410 - 0.778 |
| | DS | 0.882 | 0.649 | 0.078 | 0.571 - 0.727 | 0.887 | 0.661 | 0.080 | 0.581 - 0.741 |
| | IS | 0.701 | 0.471 | 0.112 | 0.359 - 0.583 | 0.706 | 0.476 | 0.113 | 0.363 - 0.589 |
| | SDS | 0.933 | 0.773 | 0.074 | 0.699 - 0.847 | 0.940 | 0.785 | 0.075 | 0.710 - 0.860 |
| | SIS | 0.956 | 0.634 | 0.036 | 0.598 - 0.670 | 0.963 | 0.642 | 0.037 | 0.605 - 0.679 |
| C. laurifolius | CS | 0.738 | 0.338 | 0.114 | 0.224 - 0.452 | 0.743 | 0.370 | 0.132 | 0.238 - 0.502 |
| | DS | 0.830 | 0.403 | 0.101 | 0.302 - 0.504 | 0.832 | 0.445 | 0.119 | 0.326 - 0.564 |
| | IS | 0.797 | 0.413 | 0.116 | 0.297 - 0.529 | 0.800 | 0.446 | 0.132 | 0.314 - 0.578 |
| | SDS | 0.884 | 0.594 | 0.119 | 0.475 - 0.713 | 0.887 | 0.711 | 0.182 | 0.529 - 0.893 |
| | SIS | 0.830 | 0.652 | 0.167 | 0.485 - 0.819 | 0.831 | 0.802 | 0.279 | 0.523 - 1.081 |

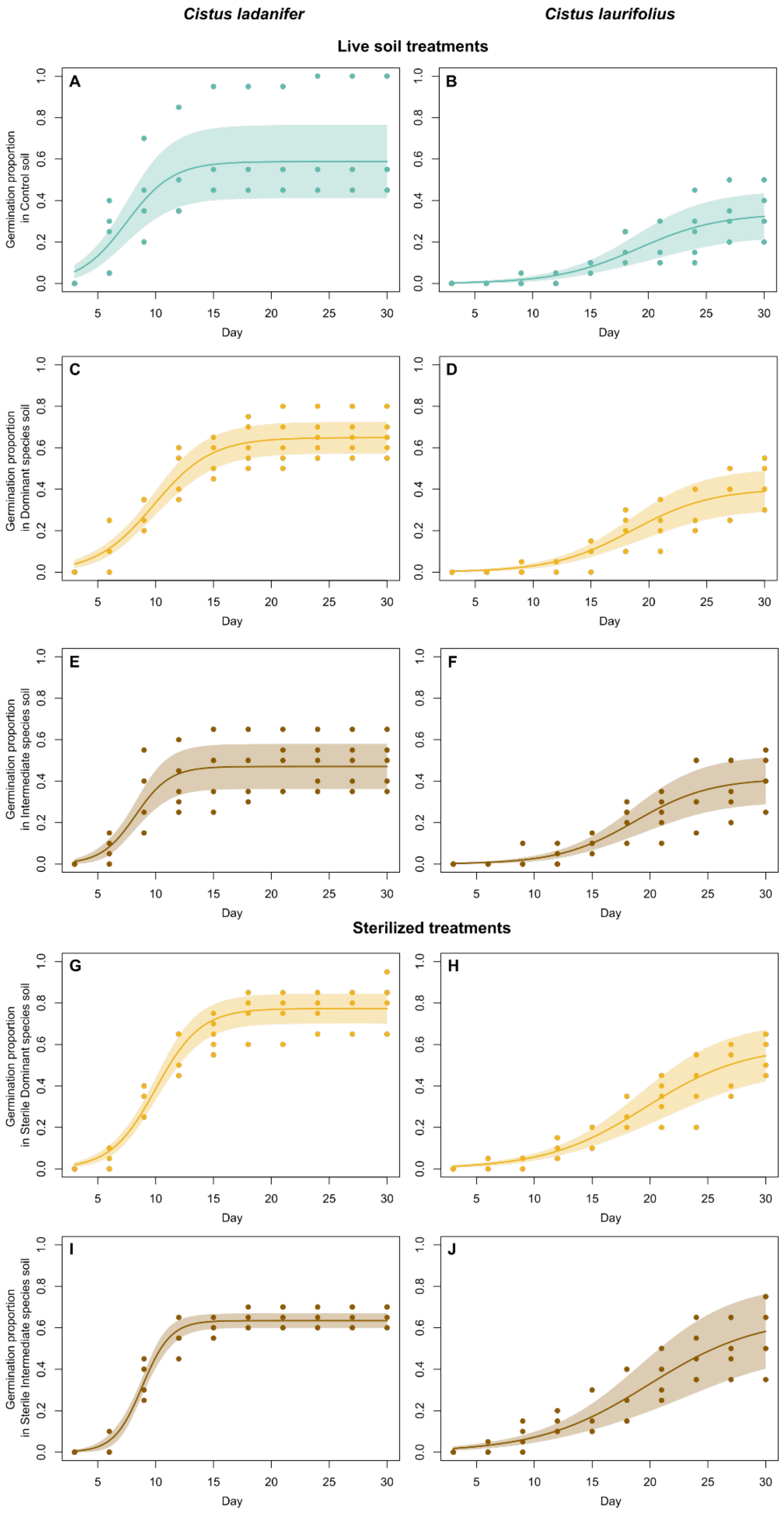

**Fig. S2. Accumulated germination proportion over time.** Panels A–F represent live soil treatments, and Panels G–J represent sterilized soil treatments. Specifically: (A–B) control soil, (C–D) dominant species' (*Cistus ladanifer*) soil, (E–F) intermediate species' (*Quercus pyrenaica*) soil, (G–H) sterilized dominant species' soil, and (I–J) sterilized intermediate species' soil. Within each pair, germination of *Cistus ladanifer* is shown on the left, and *Cistus laurifolius* on the right. Solid lines indicate the mean germination predictions from mixed-effects logistic models, while shaded areas denote the standard errors.

#### Appendix S3. Nutrient variation with distance from the intermediate species

The presence of the intermediate species significantly alters microbial communities, metabolites, and nutrients of the soils. While metabolites and microorganisms play a crucial role in nutrient cycling and availability , conducting a comprehensive analysis of their interactions is challenging due to their complexity and potential interactions. Instead, to deeply investigate how nutrient varies with distance from the tree, we conducted a detailed analysis of soil nutrient concentrations, including pH and conductivity, which influence nutrient absorption, availability and mineralization processes (Neina, 2019; Canarini *et al.*, 2019). We fitted GAMMs for each nutrient individually, including the *Q. pyrenaica* tree identity as random variable. We observed significant effects of distance on most soil properties (see Fig. S2), including Nitrogen, Phosphorus, pH, Conductivity, Nitrates, and Phosphates ($p < 0.05$). Notably, nutrient concentrations were generally highest closer to the tree, suggesting that tree-conditioned soils are enriched in nutrients. The enrichment of soil nutrients under trees can be attributed to multiple plant-mediated processes, such as litterfall deposition, root exudation, and enhanced microbial activity (Canarini *et al.*, 2019; Veen *et al.*, 2019).

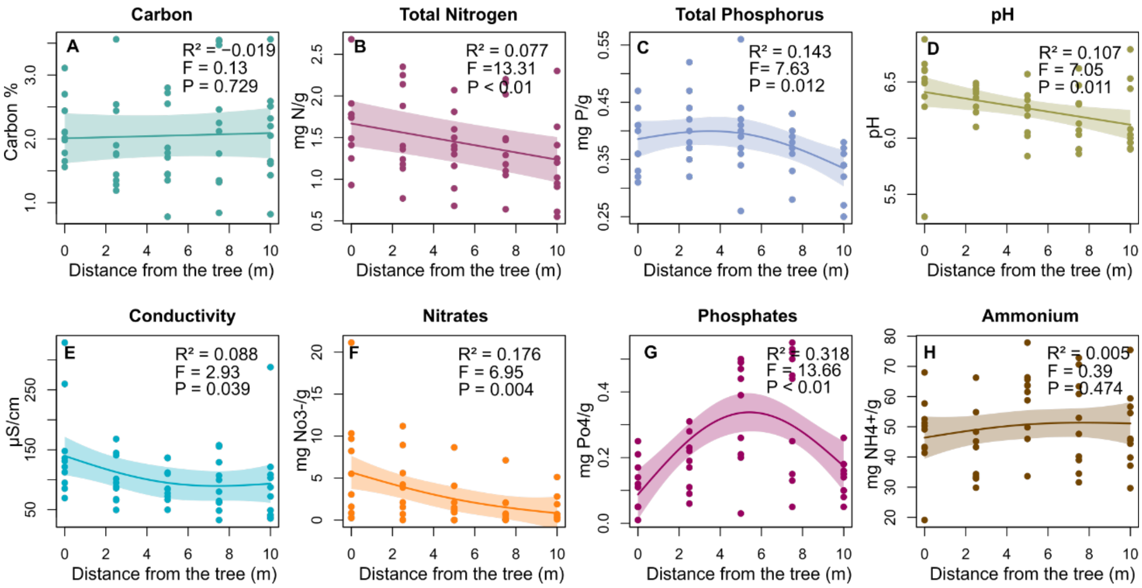

**Fig. S3. Nutrient variation with distance from the intermediate species.** Panels show the variation in different soil nutrients with distance from the *Q. pyrenaica* tree. (A) Carbon, (B) Nitrogen, (C) Phosphorus, (D) pH, (E) Conductivity, (F) Nitrates, (G) Phosphates, (H) Ammonium. Adjusted $R^2$, F statistics and p-values are also provided.

**Appendix S4. Assignment of age values to *Quercus pyrenaica* individuals**

To obtain a more accurate understanding of our study system and estimate the age of the intermediate species (*Quercus pyrenaica*) included in our research, we combined standard DBH measurements with tree-ring counts. For each of the 10 trees used in the vegetation inventory (see Methods: Field observations in the main text), DBH was measured, and a single increment core was extracted at approximately 30–40 cm above the ground using an increment borer. Cores were air-dried, mounted on wooden supports, and progressively sanded using three grades of sandpaper (80, 320, and 600 grit) until growth rings were clearly visible. Ring counts were performed by a single experienced evaluator through visual cross-dating under a stereomicroscope. The cross-dating procedure relied on the identification of pointer years, i.e. unusually narrow or wide growth rings that serve as reference markers within and among individual chronologies. This approach provided an approximate age estimate for each tree and offers a more reliable estimate of tree age than DBH-based metrics alone. Across the 10 sampled individuals, DBH ranged from 30 to 90 cm (mean = 45.9 cm), and estimated ages ranged from 29 to 52 years (mean = 36.6 years).

#### Appendix S5. Sensitivity analysis of microbial community composition

Excluding microbial taxa (ASVs) that were present in fewer than five samples, as done in the main text, may influence our results on the microbial community composition. To assess this, we conducted a sensitivity analysis without applying this taxa filtering. Including the complete microbial community confirmed that proximity to individuals of the intermediate species significantly affects both bacterial ($R^2$=3.30%; F=1.64; P < 0.01) and fungal ($R^2$=2.55%; F=1.25; P < 0.01) communities (Fig. S4). Compositional dissimilarities reached an asymptote at approximately 3.67 meters for bacteria and 3.77 meters for fungi. These findings indicate that excluding species present in fewer than five samples does not substantially alter the observed patterns (compare Fig. 3 C and D with Fig S4 A and B).

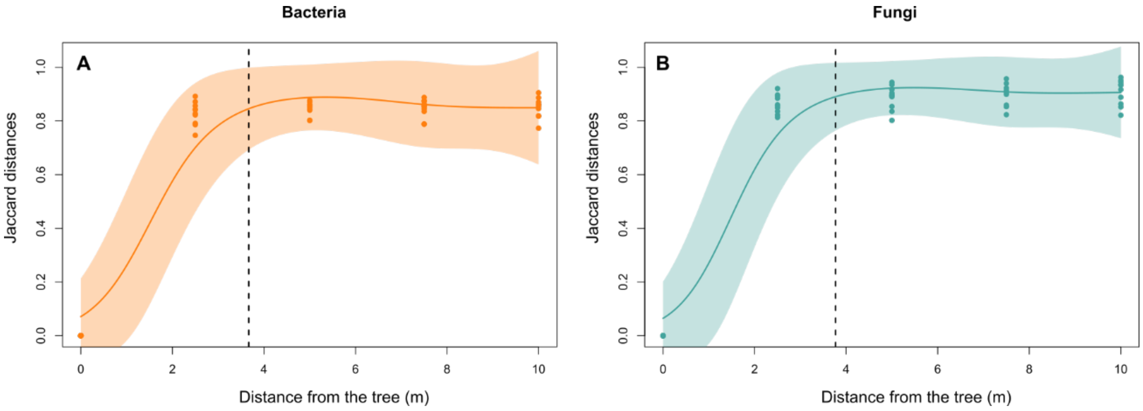

**Fig. S4. Dissimilarities in bacterial and fungal community composition as a function of distance to the tree.** Sensitivity analysis of community dissimilarities without excluding taxa occurring in fewer than five samples for (A) bacterial and (B) fungal communities. Solid lines show predictions from a generalized additive mixed model of compositional dissimilarity between the sample collected at zero distance from individuals of the intermediate species and samples collected farther away, as a function of distance. Shaded areas represent 95% confidence intervals. The vertical dotted lines indicate the inflection point in compositional dissimilarities.

**Appendix S6. Specification of simulation parameters**

This appendix provides additional details on the assumptions and parameters not described in the main text. The initial simulated starting conditions comprised 100 individuals of each competing species and 20 individuals of the intermediate species *Q. pyrenaica*, randomly distributed across a 50 × 50 grid of 1 $m^2$ cells. To explore the potential stabilizing effect of the observed soil-mediated performance shifts, we assumed that individuals of both species were otherwise ecologically identical, differing only in their experimentally observed responses across soil treatments. We assumed that each individual produce one viable seed per year. Seeds were dispersed following an exponential negative relationship with distance, favouring recruitment in nearby cells. This reflects the short-range dispersal of *Cistus* species in which most seeds fall close to the parental plant (Bastida & Talavera, 2002). The carrying capacity of each cell was set at two individuals per square meter consistent with our vegetation inventory data (mean number of individuals per $m^2$ = 2.35). Once the carrying capacity is reached, germination probability becomes zero. These simplifications act as structural constraints to prevent unbounded growth. Seed germination and survival probabilities were drawn from species-specific distributions based on experimental results. More specifically, seed germination probabilities were drawn from uniform distributions, with limits corresponding to the 95% confidence intervals estimated in the experiments for each species and soil type (Table S2). To include the effects of soil conditions on growth, we assumed that plant size influenced survival probabilities (Ps, Andivia *et al.*, 2021), following the logistic equation:

$$\mathrm{P_s}(size) = \frac{1}{1 + e^{-(x + y\, size)}}$$

where x represents the basal log-odds of survival, set at 1.386 (equivalent to a survival probability of 0.8). We established this basal survival probability based on demographic dynamics of *Cistus ladanifer*, which has a maximum lifespan of approximately 20 years (Oria-de-Rueda *et al.*, 2008). A basal survival probability of 0.8 results in only 1% of individuals reaching 20 years, approximating the observed longevity. The parameter *y* modulates the effect of plant size on survival probability, with *y*=0 maintaining a constant survival probability of 0.8 regardless of size. Higher y values increase the positive effect of plant size on survival. As the actual relationship between plant size and survival remains unknown, we conducted simulations across 21 values of *y* from 0 to 2 in 0.1 increments (Table S3). For each species and soil type, survival probabilities were drawn from a uniform distribution, constrained by the probabilities estimated from the equation above and plant sizes within the 95% mean confidence intervals obtained experimentally. We ran 25 independent simulation replicates for each scenario (21 y values X 25 replicates = 525), each spanning 100 iterations, representing years.

At the end of each simulation, we assessed the correspondence between observed and simulated abundance distributions and evaluated the population stability of both species, as detailed in the main text. We found that an average increase of 0.08 in the survival probability of *C. laurifolius* in soils conditioned by the intermediate species, relative to soils conditioned by K (y = 0.8; Fig. S5, Table S3), produced the most accurate spatial patterns while also predicting stable coexistence of the *Cistus* species.

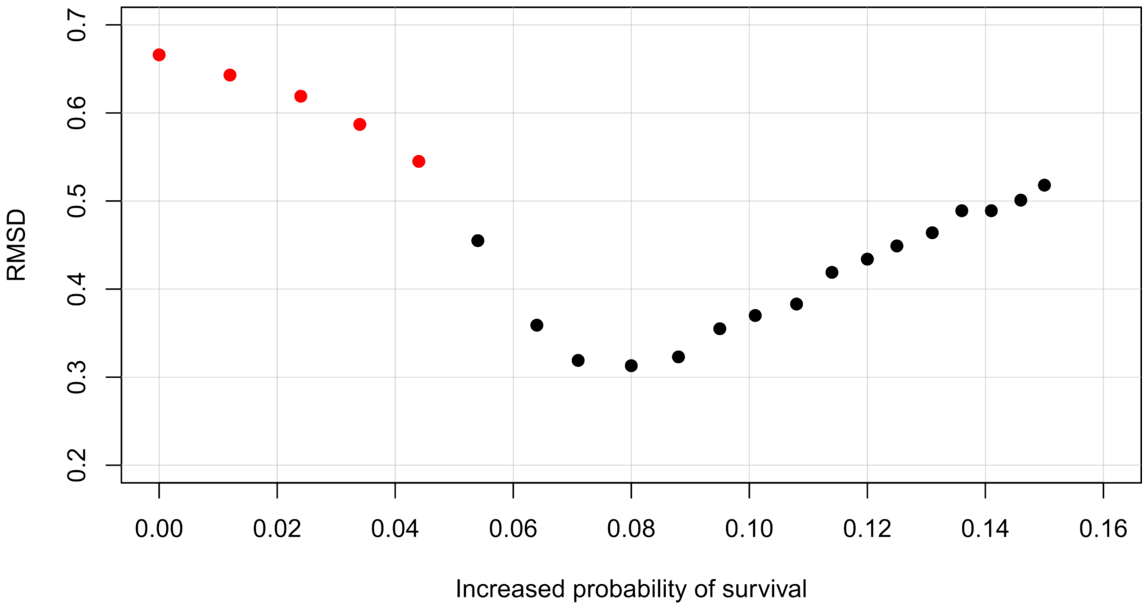

**Fig. S5. Root mean square deviation (RMSD) between simulated and observed spatial abundance patterns as a function of increased survival probability of the weak competitor in soils conditioned by the intermediate species.** Each point represents the RMSD between simulated and empirical spatial abundance patterns for a given increase in the survival probability of the weak competitor (*Cistus laurifolius*) in soils conditioned by the intermediate species (*Quercus pyrenaica*), relative to those conditioned by the dominant species (*C. ladanifer*), averaged across 25 simulation replicates (see ΔSurv.Prob. in Table S3). Lower RMSD values indicate a closer match between simulation outputs and observed field patterns. Red points denote parameter combinations that result in unstable species coexistence, while black points correspond to scenarios where stable coexistence is maintained.

**Table S3. Results of increasing the effect of size on survival probability.** Simulations were conducted by varying the effect of size on survival probability (*y*) from 0 to 2, in increments of 0.1. Results are presented for both *Cistus laurifolius* and *Cistus ladanifer*. The root mean square deviation (RMSD) quantifies the mismatch between observed and predicted abundance patterns, averaged across both species. Stability was assessed based on the statistical significance (*P*) of the relationship between abundance and time over the final 20 years of each simulation. *DS.Surv.Range* and *IS.Surv.Range* indicate the ranges of survival probabilities in soils conditioned by the dominant species and the intermediate species, respectively, for both shrub species. *ΔSurv.Prob* represents the difference between the mean survival probability in soils conditioned by the intermediate species and that in soils conditioned by the dominant species. Values shown in bold correspond to *y* = 0.8, the point at which simulated spatial patterns most closely matched the observed patterns. This value—corresponding to an increase of just 0.08 in the survival probability of *C. laurifolius* in soils conditioned by the intermediate species compared to those conditioned by the dominant species—was sufficient to promote stable coexistence in the simulated communities.

| | | C. laurifolius | | | | C. ladanifer | | | |
|---|---|---|---|---|---|---|---|---|---|
| **y** | **RMSD** | ***P* value** | **DS.Surv.Range** | **IS.Surv.Range** | **Δ Surv.Prob** | ***P* value** | **DS.Surv.Range** | **IS.Surv.Range** | **Δ Surv.Prob** |
| 0.0 | 0.666 | 0.000 | 0.800 - 0.800 | 0.800 - 0.800 | 0.000 | 0.801 | 0.800 - 0.800 | 0.800 - 0.800 | 0.000 |
| 0.1 | 0.643 | 0.000 | 0.796 - 0.806 | 0.808 - 0.817 | 0.012 | 0.857 | 0.799 - 0.804 | 0.796 - 0.802 | -0.002 |
| 0.2 | 0.619 | 0.000 | 0.791 - 0.811 | 0.815 - 0.834 | 0.024 | 0.715 | 0.797 - 0.808 | 0.793 - 0.803 | -0.005 |
| 0.3 | 0.587 | 0.000 | 0.787 - 0.817 | 0.823 - 0.849 | 0.034 | 0.680 | 0.796 - 0.812 | 0.789 - 0.805 | -0.007 |
| 0.4 | 0.545 | 0.000 | 0.782 - 0.822 | 0.830 - 0.863 | 0.044 | 0.647 | 0.795 - 0.816 | 0.785 - 0.807 | -0.009 |
| 0.5 | 0.455 | 0.214 | 0.777 - 0.827 | 0.837 - 0.876 | 0.054 | 0.585 | 0.793 - 0.819 | 0.781 - 0.808 | -0.011 |
| 0.6 | 0.359 | 0.844 | 0.773 - 0.832 | 0.844 - 0.887 | 0.064 | 0.350 | 0.792 - 0.823 | 0.777 - 0.810 | -0.015 |
| 0.7 | 0.319 | 0.760 | 0.768 - 0.837 | 0.850 - 0.898 | 0.071 | 0.315 | 0.791 - 0.827 | 0.773 - 0.811 | -0.017 |
| **0.8** | **0.313** | **0.062** | **0.763 - 0.842** | **0.856 - 0.908** | **0.080** | **0.139** | **0.789 - 0.830** | **0.769 - 0.813** | **-0.019** |
| 0.9 | 0.323 | 0.046 | 0.758 - 0.847 | 0.862 - 0.917 | 0.088 | 0.077 | 0.788 - 0.834 | 0.765 - 0.815 | -0.021 |
| 1.0 | 0.355 | 0.064 | 0.753 - 0.851 | 0.868 - 0.925 | 0.095 | 0.073 | 0.787 - 0.837 | 0.761 - 0.816 | -0.023 |
| 1.1 | 0.370 | 0.091 | 0.748 - 0.856 | 0.874 - 0.933 | 0.101 | 0.045 | 0.785 - 0.841 | 0.757 - 0.818 | -0.026 |
| 1.2 | 0.383 | 0.092 | 0.743 - 0.860 | 0.879 - 0.940 | 0.108 | 0.012 | 0.784 - 0.844 | 0.752 - 0.819 | -0.028 |
| 1.3 | 0.419 | 0.530 | 0.737 - 0.865 | 0.884 - 0.946 | 0.114 | 0.315 | 0.782 - 0.847 | 0.748 - 0.821 | -0.031 |
| 1.4 | 0.434 | 0.268 | 0.732 - 0.869 | 0.889 - 0.951 | 0.120 | 0.202 | 0.781 - 0.851 | 0.744 - 0.822 | -0.033 |
| 1.5 | 0.449 | 0.469 | 0.727 - 0.873 | 0.894 - 0.956 | 0.125 | 0.057 | 0.780 - 0.854 | 0.739 - 0.824 | -0.035 |
| 1.6 | 0.464 | 0.613 | 0.721 - 0.877 | 0.899 - 0.961 | 0.131 | 0.170 | 0.778 - 0.857 | 0.735 - 0.825 | -0.037 |
| 1.7 | 0.489 | 0.418 | 0.716 - 0.881 | 0.903 - 0.965 | 0.136 | 0.241 | 0.777 - 0.860 | 0.730 - 0.827 | -0.039 |
| 1.8 | 0.489 | 0.596 | 0.710 - 0.884 | 0.908 - 0.968 | 0.141 | 0.188 | 0.775 - 0.863 | 0.726 - 0.828 | -0.042 |
| 1.9 | 0.501 | 0.989 | 0.705 - 0.888 | 0.912 - 0.972 | 0.146 | 0.386 | 0.774 - 0.866 | 0.721 - 0.830 | -0.044 |
| 2.0 | 0.518 | 0.767 | 0.699 - 0.891 | 0.916 - 0.975 | 0.150 | 0.226 | 0.772 - 0.869 | 0.717 - 0.831 | -0.047 |

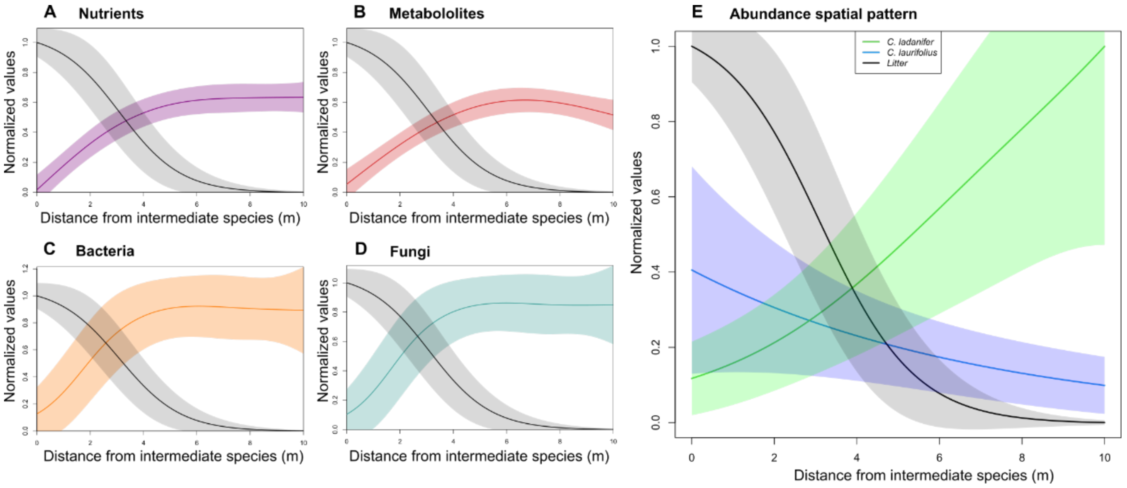

**Fig. S6. Distance-dependent shifts in soil biochemistry, microbial communities, and shrub distribution around the intermediate species, with associated litter cover.** The presence of the intermediate species (*Quercus pyrenaica*) significantly alters the composition of nutrients (A), metabolites (B), bacteria (C), fungi (D), and the spatial distribution of the competing shrub species (E) (see Fig. 3). Solid lines show predictions from generalized additive mixed models of compositional dissimilarity between samples collected at zero distance from individuals of the intermediate species and those collected farther away, as a function of distance. Shaded areas represent 95% confidence intervals. Litter cover was measured as the percentage of ground surface covered by leaf litter in each 1 $m^2$ plot, simultaneously with counts of *Cistus ladanifer* and *C. laurifolius* individuals along 10 × 1 m transects (see Methods: Field observation). Values are normalized to the maximum observed within each panel. Litter cover is shown in black, with shaded areas representing 95% confidence intervals. Predictions for litter cover are based on a generalized additive mixed model with a binomial error distribution and a logit link function ($R^2$ = 65.7%; $F$ = 23.32; $P$ < 0.01). These results indicate that the areas where compositional changes occur are largely covered by tree litter.

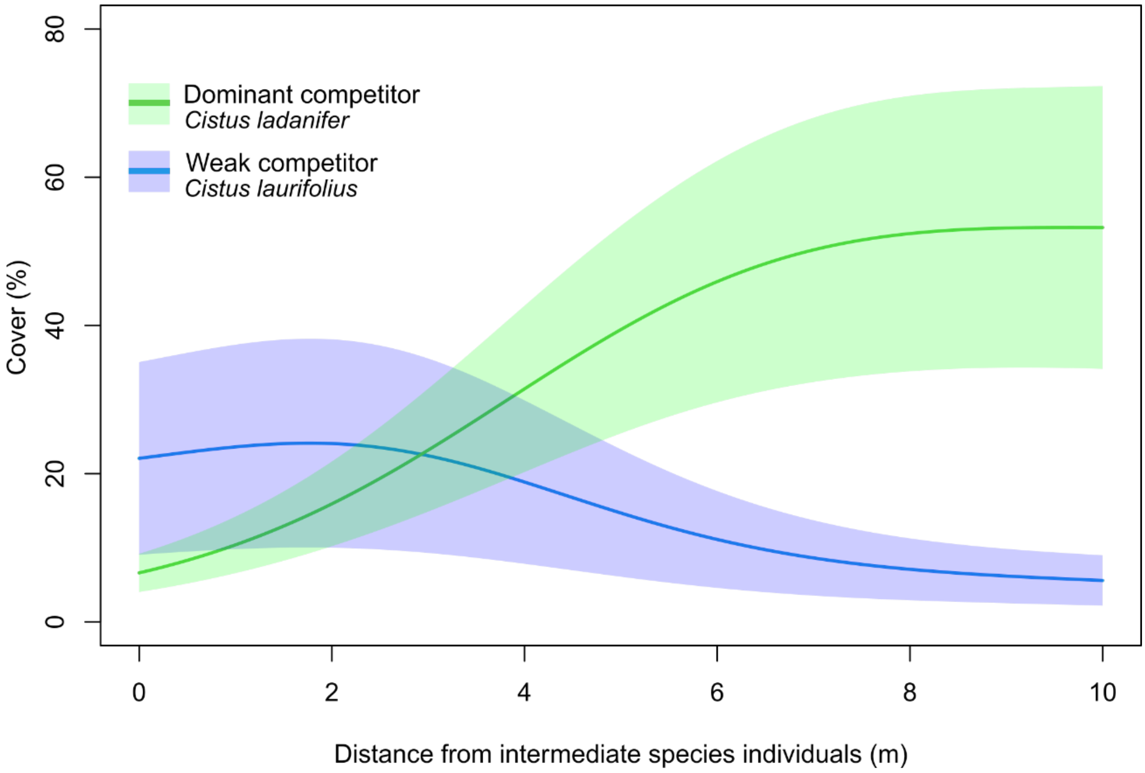

**Fig. S7. The presence of the intermediate species alters the spatial distribution of the competing shrub species.** Predicted spatial distribution of the cover (per 1 $m^2$) of the dominant (green; $R^2$=29.3%, F=280.1, $P$ < 0.01) and the weak (blue; $R^2$=7.64%, $F$=150.7, $P$< 0.01) competitor species as a function of distance from individuals of the intermediate species. Solid lines show the fitted values from generalized additive mixed models (GAMMs), with shaded areas indicating 95% confidence intervals. GAMMs were fitted using a Poisson error distribution and a log link function, and included as a random the identity of *Quercus pyrenaica* individuals.
